# *De novo* generation of antibody CDRH3 with a pre-trained generative large language model

**DOI:** 10.1101/2023.10.17.562827

**Authors:** Haohuai He, Bing He, Lei Guan, Yu Zhao, Guanxing Chen, Qingge Zhu, Calvin Yu-Chian Chen, Ting Li, Jianhua Yao

## Abstract

Artificial Intelligence (AI) techniques have made great advances in assisting antibody design. However, antibody design still heavily relies on isolating antigen-specific antibodies from serum, which is a resource-intensive and time-consuming process. To address this issue, we propose a Pre-trained Antibody generative large Language Model (PALM) for the de novo generation of artificial antibodies heavy chain complementarity-determining region 3 (CDRH3) with desired antigen-binding specificity, reducing the reliance on natural antibodies. We also build a high-precision model antigen-antibody binder (A2binder) that pairs antigen epitope sequences with antibody sequences to predict binding specificity and affinity. PALM-generated antibodies exhibit binding ability to SARS-CoV-2 antigens, including the emerging XBB variant, as confirmed through *in-silico* analysis and *in-vitro* assays. The *in-vitro* assays validated that PALM-generated antibodies achieve high binding affinity and potent neutralization capability against both wild-type and XBB spike proteins of SARS-CoV-2. Meanwhile, A2binder demonstrated exceptional predictive performance on binding specificity for various epitopes and variants. Furthermore, by incorporating the attention mechanism into the PALM model, we have improved its interpretability, providing crucial insights into the fundamental principles of antibody design.

## Introduction

Antibody drugs, also known as monoclonal antibodies, play a vital role in biotherapy^1,2^. By mimicking the actions of the immune system, these drugs selectively target pathogenic agents such as viruses and cancer cells^3^. Compared to traditional treatments, antibody drugs offer a more specific and efficacious approach^4^. Antibody drugs have demonstrated positive outcomes in the treatment of numerous diseases^5^, including COVID-19^6^.

Developing antibody drugs is a complex process that involves isolating antibodies from animal sources, humanizing them, and optimizing their affinity^7,8^. Moreover, with advancements in technology and research methodologies, several faster and more contemporary approaches have emerged. These include isolating antibodies directly from patients^9^, immunizing transgenic mice with human immune systems^10^, and utilizing *in-vitro* discovery methods from donor and synthetic libraries^11^. The development of computational methods such as Rosetta^12^, SnugDock^13^, and artificial intelligence (AI) algorithms^14^ has greatly facilitated the prediction of antibody affinity and the optimization of the antibody design process, making it faster and more efficient. These methods have the potential to significantly accelerate the design of antibody drugs. Despite these advances, the development of antibody drugs still heavily relies on natural antibodies.

The sequence data of protein can be regarded as a language, so the large-scale pre-training models in the natural language processing (NLP) field have been used to learn the characterization pattern of the protein. Verkuil et al.^15^ employed EMS2^16^, a large language model trained only on protein sequences, to learn the deep grammar of protein language. Nijkamp et al.^17^ introduced a suite of protein language models that show state-of-the-art performance in many downstream tasks. Madani et al. proposed ProGen^18^, a method for generating protein sequences that are controlled by protein properties. However, the generation of antibodies with high affinity to specific antigen epitopes remains a challenging task due to the high diversity of antibodies and the scarcity of available antigen-antibody pairing data.

To address the aforementioned challenges, we propose a Pre-trained Antibody generative large Language Model (PALM) for optimizing and generating the heavy chain complementarity-determining region 3 (CDRH3), which plays a vital role in the specificity and diversity of antibodies^19,20^. To avoid the problem of lacking paired datasets and improve model performance, we pre-train the Roformer^21^ on a large number of unpaired protein sequences, followed by fine-tuning and evaluating the model on an antigen-antibody affinity dataset. Subsequently, we use the fine-tuned model and antigen-antibody pairing data to generate the CDRH3 of the antibody.

To evaluate the affinity of the antibodies generated by PALM for antigens, we utilized a combination of antigen-antibody docking^13^ and AI-based methods. Previous AI methods have utilized antibody structural information to predict the antigen-antibody binding affinity^22–24^. However, collecting confident protein structures through wet-lab experiments is a time-consuming and labor-intensive process^25^. Some other methods utilize antibody sequences, which are relatively cheaper and easier to obtain, to predict affinity for specific antibodies^26,27^. Although such methods employ large-scale pre-trained language models to achieve even more accurate affinity predictions^28,29^, their prediction ability is limited to the trained antigens since they do not consider the antigen information. When dealing with unknown antigens, such as in new mutation affinity tasks like XBB, the lack of antigen information is a significant limitation. To address this issue, we developed the A2binder for evaluating antibody-antigen affinity. A2binder uses a large-scale pre-trained model for sequence feature extraction from both antigens and antibodies, followed by feature fusion and final affinity prediction using MF-CNN. This approach enables A2binder to achieve accurate and generalizable affinity predictions even for unknown antigens.

In this study, we introduced two novel methods, PALM and A2Binder, and developed a comprehensive workflow for antibody generation and affinity optimization. With the abundance of antigen-antibody data generated during the COVID-19 pandemic, we evaluated our approach using COVID-19 data. Our results demonstrated the successful generation of antibodies targeting a stable peptide in the HR2 region of COVID-19 and antibodies with higher affinity for the new variant XBB. In conclusion, we have established an artificial intelligence framework for antibody generation and evaluation, which has the potential to significantly accelerate the development of antibody drugs.

## Results

### The framework of PALM and A2Binder

The workflow and model framework of the pre-trained Antibody generative large Language Model (PALM) and the antigen-antibody binder (A2binder) are shown in Figure 1. The purpose of PALM is to generate the CDRH3 sequence in the antibody. As illustrated in Figure 1a, the CDRH3 region plays the most vital role in determining an antibody’s binding specificity against a particular antigen sequence. As illustrated in Figures 1b, 1c, and 1d, PALM is a transformer-like model ^30^ that uses the ESM2^16^-based Antigen model as the encoder and Antibody Roformer as the decoder. The training of PALM and A2binder consists of three steps: first, we pre-train two language models on unpaired antibody heavy and light chain sequences, respectively. Then we construct A2binder and fine-tune it using paired affinity data. Finally, we construct PALM using the pre-trained ESM2^16^ and Roformer models and train it on paired Antigen-CDRH3 data for designing and evaluating the AI-generated CDRH3. The data statistics used for training can be found in Supplementary Table S1, while the details of the training and model hyperparameter settings can be found in Supplementary Section ‘Training detail’ and Table S2.

**Figure 1.**
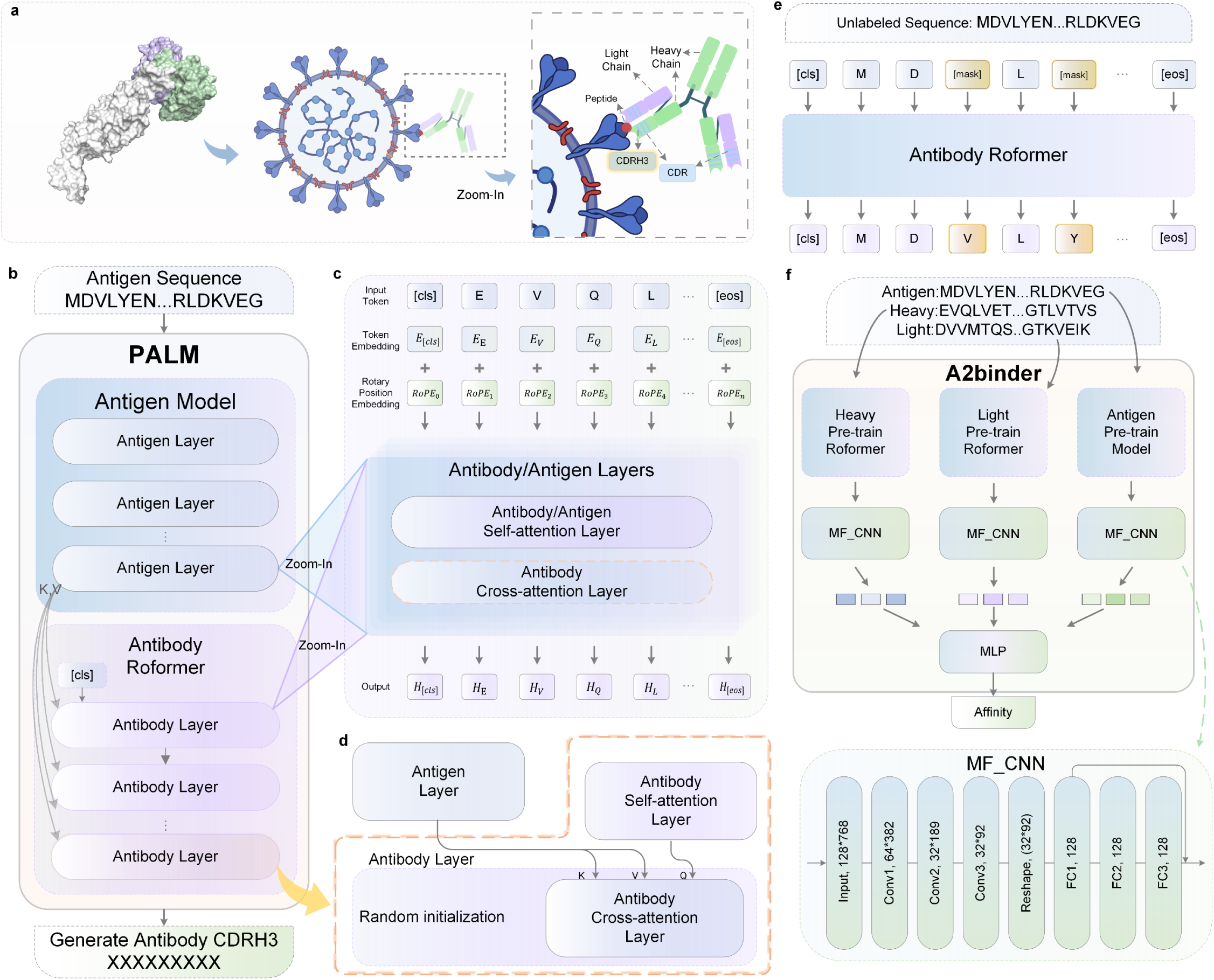
Overview of the PALM and A2binder workflow. **a**, Schematic of an antibody binding to the epitope region of an antigen. The antibody paratope comprised of CDRs from the heavy and light chains mediates the recognition and binding of the antigen. The CDRH3 loop, as the third CDR of the antibody heavy chain, plays an essential role in enabling specific antigen binding. **b**, The framework of PALM. It’s a Transformer-like neural network containing an antigen encoder model and an antibody decoder model. It takes the antigen sequence as input and generates a CDRH3 antibody sequence aiming to bind to the input antigen. The antigen encoder model is a ESM2-based model, which comprised of 12 antigen layers that were pre-trained using UniRef50 protein sequences and fine-tuned using antigen sequences. The antibody decoder is a RoFormer –based model, which contains 12 antibody layers that were pre-trained and fine-tuned using antibody sequenses. The key (K) and value (V) matrices from the last antigen layer are passed to every antibody layer as the input of the cross-attention sub-layer. **c**, Internal architecture of the antigen layer and antibody layer. Both antigen layer and antibody layer have two basic sub-layers, including a fully connected feed-forward sub-layer and a multi-head self-attention sub-layer. Additionally, the antibody layer uniquely includes cross-attention sub-layers. Input tokens of each layer are represented by the sum of token embeddings and rotary position embeddings, while the output is a high-dimensional vector representation for each input token. **d**, The cross-attention sub-layer is the key to combine the high-dimensional representation of antigen sequence (K and V matrices) and in-context antibody sequence(Q matrix). **e**, Schematic of the self-supervised pre-training of antibody RoFormer. Unpaired antibody sequences were used to pre-train the antibody RoFormer via masked language modeling. Sequences were tokenized and had partial random tokens masked. The model was trained to predict the identity of the masked tokens, learning generalizable representations of antibody sequences. **f**, The framework of A2Binder. It takes the antigen sequence along with antibody heavy and light chain sequences as input. Each sequence is encoded by passing through a pre-trained encoder and a Multi-Fusion Convolutional Neural Network (MF-CNN) feature extractor. The MLP (a multilayer perceptron) model finally fuses the features from all three sequences to predict antibody-antigen binding affinity. The architecture of the MF-CNN is shown below, comprising 3 convolutional layers and 3 fully-connected layers. The encoder for both the antibody heavy chain and light chain are built upon the pre-trained antibody RoFormer, whereas the antigen encoder utilizes the ESM2.

To pre-train the antibody Roformer, the characterization patterns of both light and heavy chain sequences of antibodies were utilized in developing a feature extraction pre-training model for COVID-19 antibodies. As illustrated in Figure 1e, the model’s architecture was based on the Roformer^21^ and was trained using the pre-training strategy of learning language representation patterns. The OAS database^31,32^ contains over 1 billion unpaired light and heavy chain sequences.

Our objective was to uncover generalizable patterns about SARS-CoV-2 antibody sequences, so we screened this repository to extract 81,750,886 heavy chain and 17,754,502 light chain sequences from COVID-19 patient data. While derived from COVID-19 immune repertoires, these sequences likely only show limited SARS-CoV-2 specificity and contain many off-target binders not related to COVID-19. However, analyzing this large-scale sequence corpus from COVID-19 patients enabled our model to learn structural patterns common across diverse antibodies beyond any single antigen specificity. The Mask Amino Acid (MAA) task was then applied to obtain neural network models to characterize patterns of antibody light and heavy chain sequences.

We fine-tuned A2binder on the antigen-antibody affinity task to enable it to learn the rules of antigen-antibody binding. The architecture of the A2binder is shown in Figure 1f, it encompasses of pre-trained language models for feature extraction, serving to extract information from light chain, heavy chain, and antigen sequences. Following each language model is a multi-layered CNN architecture named Multi-Fusion CNN (MF-CNN). The light-chain and heavy-chain Roformers from pre-training are used to extract information about light and heavy chains. We also employ a large-scale pre-trained model ESM2^16^ for extracting features of antigen sequences. The MF-CNN was designed to combine the sequence feature extraction outputs from pre-training models. The output from the concatenation of the features from MF-CNN was utilized to predict the affinity. Further introduction to the model can be found in the ‘Methods’ section.

Following previous work^33^, we used the pre-trained ESM2 model as our antigen model and the pre-trained heavy Roformer from step one as our decoder to create PALM. As illustrated in Figures 1b and 1d, the antigen and antibody models are stacked by multiple antigen and antibody layers, respectively. PALM incorporates encoder and decoder self-attention layers, with their initial weights inherited from the pre-trained ESM2 model and antibody Roformer, respectively. The decoder also includes an Antibody Cross-attention Layer, which is randomly initialized and fine-tuned using paired CDRH3-antigen sequence data for the sequence-to-sequence task. The last antigen layer passes k, v matrices into all Antibody Cross-attention Layers, while the q matrix comes from the Antibody Self-attention Layer. Through the attention mechanism^30^, PALM realizes the translation task from antigen to CDRH3.

### Pre-training allows the model to learn a better representation of antibodies

During the pre-training stage, the model learns potential representation patterns of antibody sequences through exposure to a diverse range of antibody sequences, facilitating effective feature extraction from the input antibody sequences. The prediction accuracy was 92.74% and 94.14% for heavy chain Roformer and light chain Roformer, respectively, indicating excellent pattern characterization capability of the pre-trained model.

We further investigated whether the pre-trained model could differentiate the antigenic region, type, and binding affinity targeted by the antibody. Initially, we utilize the CoV-AbDab^34^ database, which contains variant and epitope information of the antigens. From this database, we filter out antibody data that specifically target a single variant of the novel coronavirus. Subsequently, we input the antibody sequences into both the untrained Roformer and the pre-trained Roformer to obtain the embedding of the antibody sequence. We ran t-SNE to visualize the feature distribution, as shown in Figure 2a. The feature representation of the untrained Roformer was scattered, whereas the pre-trained Roformers exhibited feature clustering according to the same antigen.

**Figure 2.**
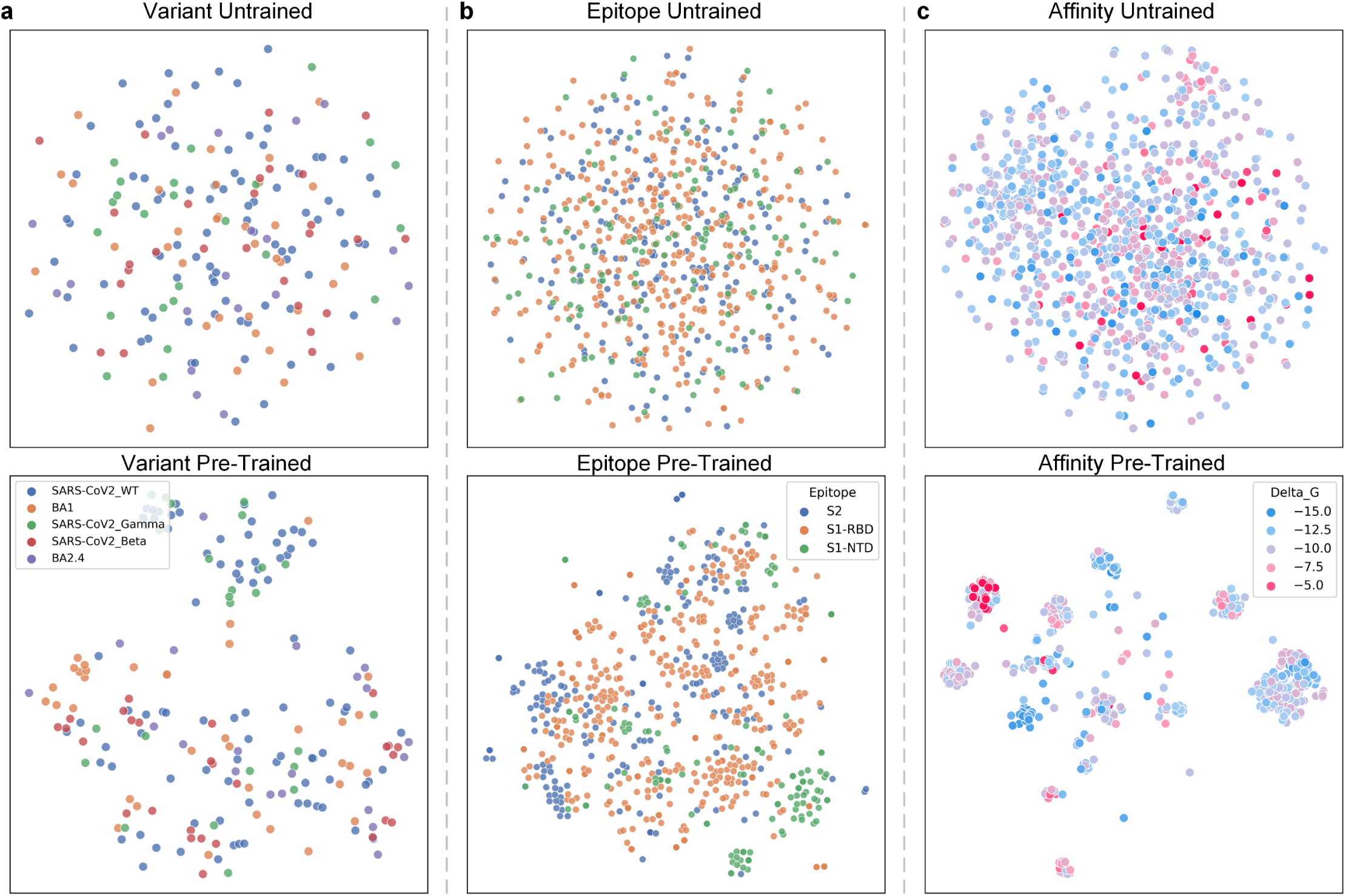
Comparison of latent capabilities between pre-trained and untrained models. Dimensionality reduction visualizations demonstrate the pre-trained model’s inherent ability to capture the feature of antibodies that determine the antigen binding specificity and affinity. **a**, T-SNE projection of sequence embeddings for antibodies selectively targeting distinct SARS-CoV-2 variants. Antibodies that bound to multiple variants were eliminated. **b**, T-SNE projection of model embeddings for antibodies specifically targeting unique epitopes of SARS-CoV-2. Antibodies that bound to multiple epitopes were eliminated. Antibody sequences used in subgraph a and b are from CoV-AbDab dataset. **c**, T-SNE projection of model embeddings of antibody sequences with different binding affinity. Each point represents a single antibody sequence from the BioMap dataset, with colors indicating the binding affinity, expressed as binding free energy (ΔG).

Then, we evaluated the pre-trained model’s ability to represent epitopes. We selected antibody data from the CoV-AbDab^34^ database that target a specific epitope. Similarly, we utilized both the untrained and pre-trained models to obtain embeddings and subsequently employed t-SNE for dimensional reduction visualization. Figure 2b depicts the reduced dimensional visualization of the pre-trained model’s epitope embeddings for antibody-bound antigens, demonstrating that the pre-trained model produces aggregated embeddings for each epitope. In contrast, the model without pre-training generated scattered embeddings.

Moreover, we aimed to assess the model’s proficiency in characterizing binding abilities. Thus, we utilized antibodies from the BioMap ^35^ dataset, which includes binding free energy (Delta G) as the antigen-antibody affinity data, to visualize the dimensionality reduction of embeddings. All samples in the BioMap dataset are derived from the Biomap company. Figure 2c presents the embedding results of the model before and after pre-training. The pre-trained model effectively aggregated high and low-affinity embeddings.

Collectively, these three comparison results demonstrate that pre-training enhances the model’s ability to extract critical information, such as the antibody’s binding antigen type, region, and affinity.

### A2binder can accurately predict the antigen-antibody binding probability

The performance of A2binder was evaluated by comparing its ability to predict affinity to that of several baseline methods including ESM-F, Ens-Grad, and Vanilla BERT on multiple affinity datasets. Section “Baseline” provides detailed information about the baseline methods.

Initially, the CoV-AbDab dataset was used ^34^, resulting in 27,324 antigen-antibody pair data for 22 SARS-CoV-2 variants as antigens. Since this dataset does not contain specific affinity values, we used neutralization or non-neutralization as a label to evaluate the performance of the A2binder in a binary classification task. The details of data processing procedures are expounded upon in the dataset subsection of the Methods section.

Figures 3a, 3b, and Supplementary Table S3 illustrate the performance of A2binder and the baseline methods on the CoV-AbDab dataset. A2binder outperformed all the baseline models in terms of the area under the receiver operating characteristic (ROC-AUC) and the precision-recall area under the curve (PR-AUC). A2binder achieved a ROC of 0.930 which was a 2% performance improvement. It is observed that the BERT model without pre-training performed the worst, which highlights the importance of pre-training in obtaining the characterization of antibody and antigen sequences for model performance. We also compared the model performance under different epitopes and variants, and the results are shown in Figures 3c and 3d. The model can achieve good performance under different epitopes and variants.

**Figure 3.**
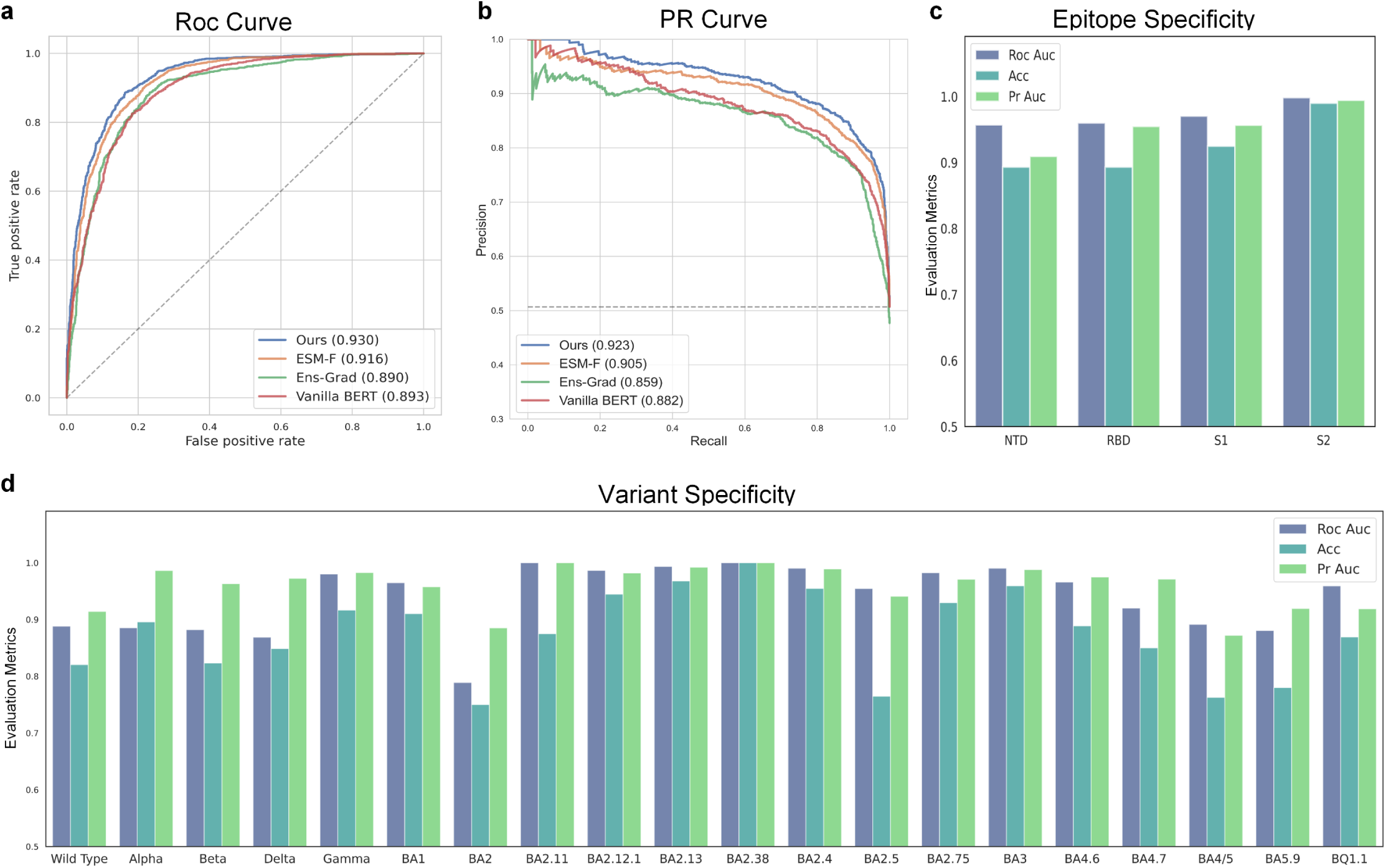
Performance comparison of A2Binder versus baseline methods for antibody-antigen binding specificity prediction. **a-b**, Receiver operating characteristic (ROC) curve (a), and precision-recall (PR) curve (b) evaluating the overall predictive performance of antibody binding specificity. Models compared include A2Binder (blue), ESM-F (orange), Ens-Grad (green), and Vanilla BERT (red). **c-d**, Performance breakdown of A2Binder in predicting antibody binding specificity by antigen epitope region (c) and variant (d). The x-axis labels indicate the different epitope categories (c) and variants (d). The CoV-AbDab dataset was split into training (80%), validation (10%) and test (10%) sets. Results shown in this comparison are based on the test set.

### A2binder can accurately predict antigen-antibody affinity

The task of predicting binding affinity values through regression is more challenging than the binary task of predicting neutralization or non-neutralization. To assess the model’s performance in predicting affinity values, we also utilized two datasets, 14H and 14L ^36^, that contain labels for affinity values. Both datasets contain a measure of the affinity of the antibody to a stable peptide in the HR2 region of SARS-CoV-2 ^37^. The heavy chains of the 14H dataset vary, while the light chain is constant, whereas the 14L dataset is the opposite. Therefore, for the 14H dataset, we used the pre-trained heavy chain Roformer to extract features from the CDRH1, 2, and 3 regions, while the 14L dataset used the pre-trained light chain Roformer. The details of the specific data processing process are in the dataset subsection of the Methods section. Figures 4a, 4b, and Supplementary Table S4 illustrate the performance comparison of models on 14H and 14L. A2binder outperformed all baseline models in Spearman’s rank correlation coefficient metrics. A2binder achieved a Spearman of 0.553 on the 14H dataset (7% improvement), and 0.688 (1% improvement) on the 14L dataset. The pre-trained model ESM-F outperformed other baseline methods in all metrics, further verifying the significance of pre-training. Additionally, the sequence pre-training of SARS-CoV-2 antigens may assist the model in learning the characterization of SARS-CoV-2-related antibody sequences more effectively.

**Figure 4.**
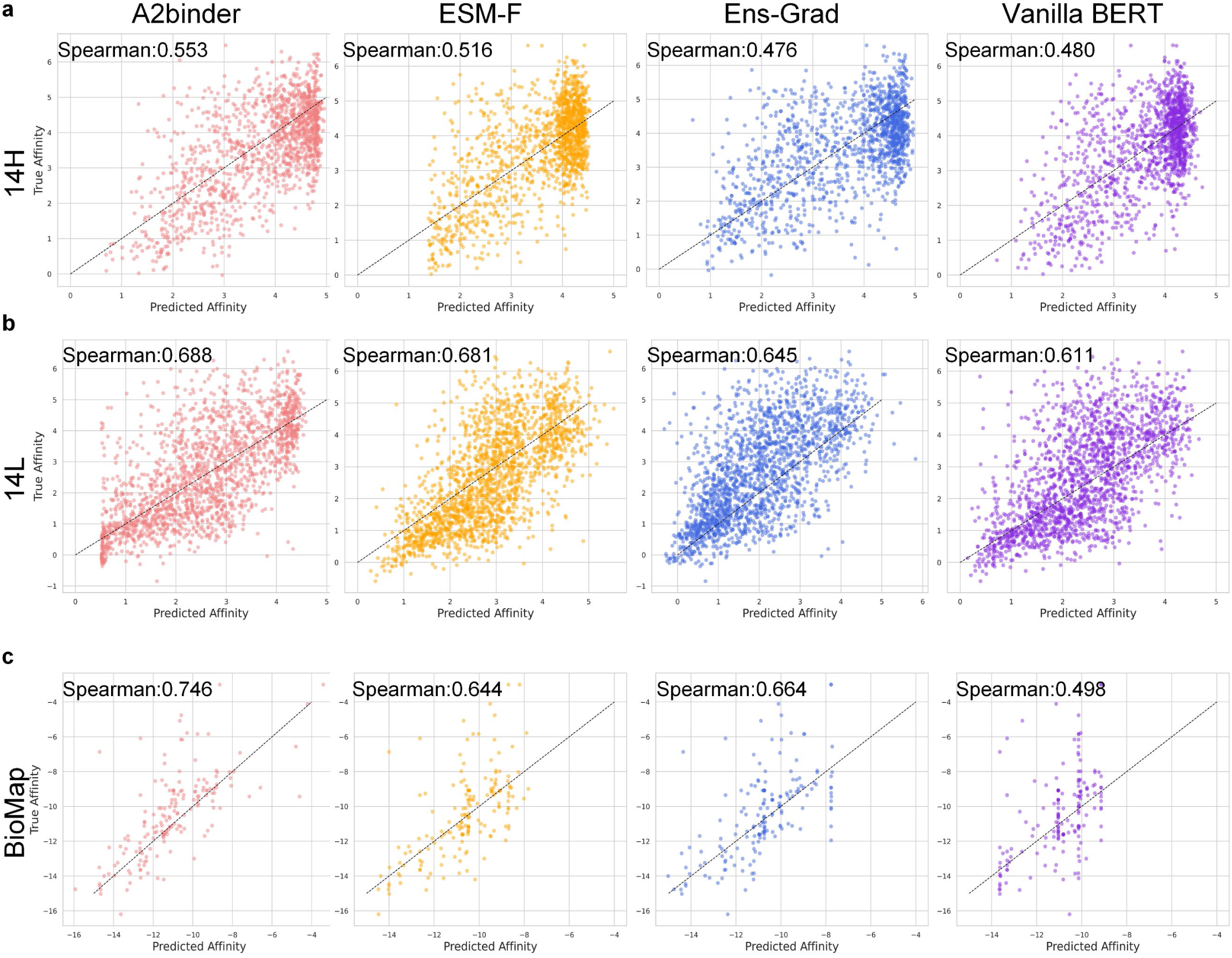
Performance comparison of A2Binder versus baseline methods for antibody-antigen binding affinity prediction. **a-c** Scatter plots showing predicted vs. true binding affinity for each model using the 14H (a), 14L (b), and BioMAP (c) datasets, respectively. Models compared include A2Binder (red), ESM-F (orange), Ens-Grad (blue), and Vanilla BERT (purple). The 14H, 14L, and BioMap datasets were each divided into training (80%), validation (10%), and test (10%) sets. Results shown in this comparison are based on the test sets. Upon statistical testing, we found A2Binder’s improvements in Spearman correlation over baselines to be significant (p<0.05 on 14H and 14L; p<0.01 on BioMap, t-test).

To verify the model’s ability to predict antibody affinities for antigens other than SARS-CoV-2, the BioMap dataset was used to evaluate the model’s prediction performance.

Figure 4c illustrates the performance comparison of the proposed model on the BioMap dataset. A2binder achieves a 6% performance improvement in reaching a Spearman of 0.746. Consistent with previous results, A2binder outperforms the baseline methods on all metrics. This further supports the model’s ability to accurately predict antigen-antibody affinity, regardless of whether the antigen is related to SARS-CoV-2 or not. This may be attributed to the use of MF-CNN architecture in A2binder, which enables the extraction of global feature output from a large-scale pre-trained model. And it is noteworthy that our model was pre-trained on a dataset that included a substantial amount of data related to COVID-19 antibodies, forming a strong foundation for predicting binding affinities for this specific antigen. However, the 14H and 14L datasets, characterized by limit ed variability in antibody sequences, present a more challenging prediction task. This may be the reason why the performance of the model in Biomap is higher than that of 14H and 14L.

### PALM can generate antibodies that are dissimilar to natural ones in sequence yet exhibit a high binding probability

To investigate the diversity of antibodies generated by the PALM, we selected the CDRH3 sequences of natural antibodies targeting the wild-type SARS-CoV-2 RBD region from the CoV-AbDab. Subsequently, we employed PALM to generate CDRH3 sequences targeting the same epitope. And we created a sequence logo plot for both artificial and natural antibodies. Figure 5a illustrates that the first three amino acids of the generated antibodies are similar to the natural antibodies since ‘ARD’ has the highest probability. The artificial antibodies exhibit greater diversity in their tail sequences, with the most probable tail being ‘DY’. Additionally, the middle regions of the generated antibodies display considerable diversity.

**Figure 5.**
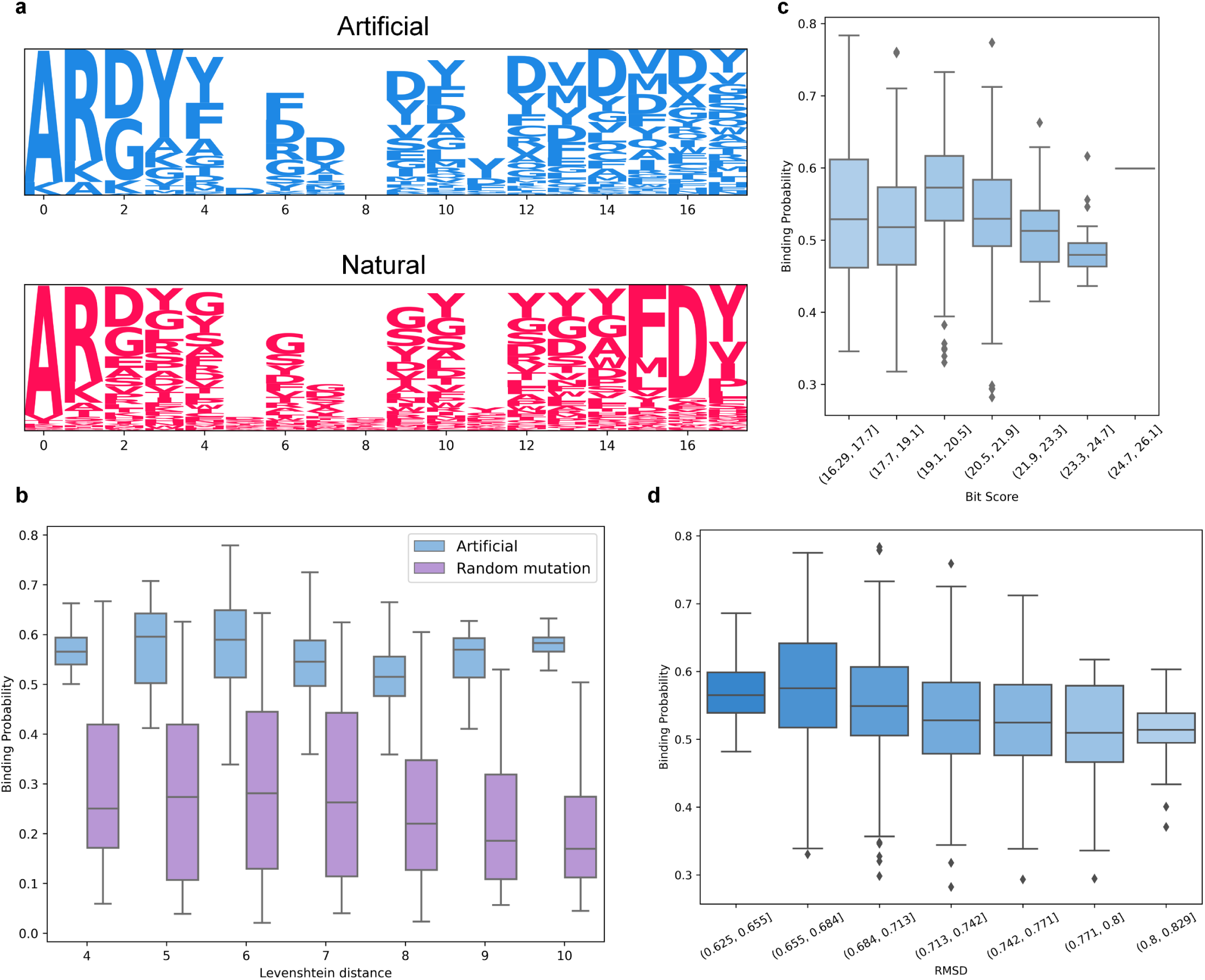
Similarity analysis of artificial and natural antibodies. **a**, Sequence logo of the CDRH3 region in artificial and natural antibodies. The CDRH3 sequences of natural antibodies are sourced from antibodies in the CoV-AbDab dataset that bind to the RBD region of wild-type SARS-CoV-2, while artificial antibodies CDRH3 sequences are obtained by inputting the RBD sequence of the wild-type SARS-CoV-2 to the PALM. **b**, Comparison of A2binder-predicted binding probabilities to the wild-type SARS-CoV-2 RBD region between generated antibodies and randomly mutated antibodies. Artificial antibodies and randomly mutated antibodies with the same Levenshtein distance to natural antibodies are compared. Boxplot showing distribution of A2binder-predicted binding probabilities across different Levenshtein distances. The x-axis denotes Levenshtein distance and the y-axis shows predicted binding probability. Blue boxes represent human engineered antibodies while purple boxes denote randomly mutated antibodies. The top whisker, top of the box, middle line, bottom of the box, and bottom whisker indicate the maximum, 75th percentile, median, 25th percentile, and minimum values, respectively. **c**, A2binder-predicted binding probabilities of artificial antibodies at different BitScore ranges. The BitScore measures the sequence similarity between artificial antibodies and natural antibodies binding to the same epitope. The x-axis denotes Bit score ranges and the y-axis shows predicted binding probability. The depth of the color indicates an increase in BitScore. The diamond represents outliers. **d**, A2binder-predicted binding probabilities of artificial antibodies at different RMSD (root mean square deviation) ranges. The RMSD measures the structure similarity between artificial antibodies and natural antibodies binding to the same epitope. The x-axis denotes RMSD ranges and the y-axis shows predicted binding probability. The depth of the color indicates an increase in RMSD value. The diamond represents outliers.

To investigate whether dissimilar sequences result in reduced binding probability, we computed the edit distance between generated antibody sequences and natural antibodies. We divided the dataset based on edit distance and employed the A2binder to predict the binding probability, as shown in Figure 5b. For comparison, we also generated sequences with random mutations and randomly generated sequences in line with the edit distance.

The results indicated that the generated antibodies exhibited a higher binding probability and did not exhibit a declining trend in probability as the edit distance increased. In contrast, the random mutation results showed a decrease in affinity probability as the edit distance increased.

Furthermore, we obtained the BitScore of the artificial antibody by Blast. A larger BitScore value indicates a higher similarity with the natural antibody. As shown in Figure 5c, the artificial antibodies did not exhibit a decrease in binding probability due to low similarity, which is consistent with the previous analysis.

To investigate the influence of structure on binding probability, we utilized AlphaFold2 (AF2) ^38^ to generate the structure of the artificial antibody and computed the Rmsd between the artificial and natural antibodies.

As depicted ed in Figure 5d, an increase in Rmsd results in a decrease in the average probability of antibody binding. This may suggest that a decrease in structural similarity could lead to a reduction in the likelihood of antigen-antibody binding. However, even in the interval with the highest RMSD, the binding probability remains higher than 0.5. In conclusion, the PALM is capable of generating a diverse set of antibody sequences with low sequence similarity, yet still exhibiting high binding probabilities.

### PALM can generate antibodies that bind to the desired epitopes on the antigens

To determine if the sequences generated by the PALM can target the correct site, we generated antibodies against various domains. As illustrated in Figure 6a, the antibodies generated by the PALM can be similar to the natural antibodies that target the same domain.

**Figure 6.**
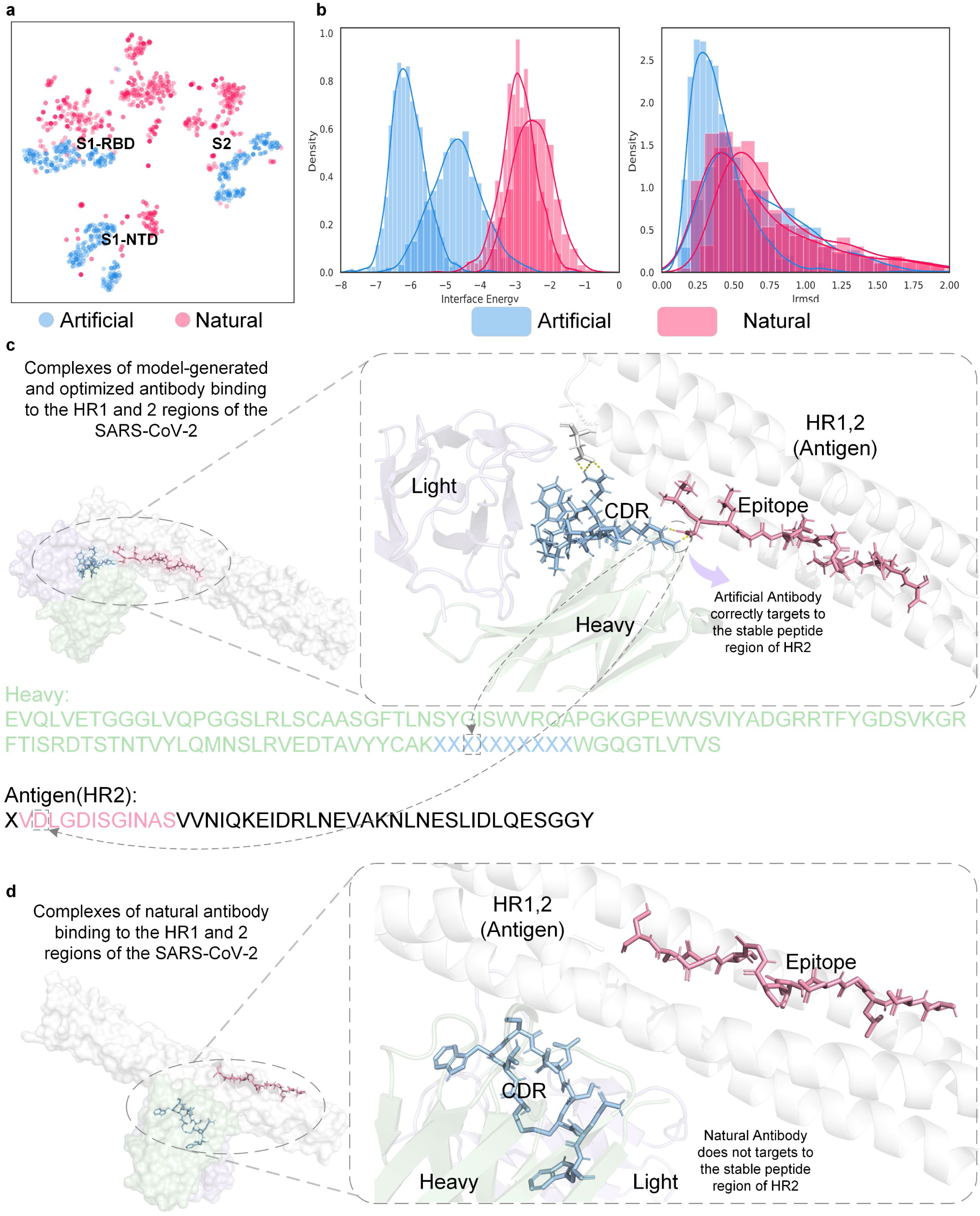
The performance of PALM in generating artificial antibodies binding to the different epitopes of SARS-CoV-2. **a**, The t-SNE plot shows that artificial and natural antibodies with identical binding specificity are clustered in the same group. Each blue dot indicates an artificial antibody, while a red dot indicates a natural antibody. Each antibody sequence is embedded using the pre-trained light chain and heavy chain Roformers, followed by dimensionality reduction to obtain feature representations using t-SNE. **b**, Density distribution plots of the binding affinity of the artificial and natural antibodies targeting the same epitope. The binding affinity represented by Interface Energy and Irmsd are derived from 1000 optimized poses of the antibody-antigen binding complex using SnugDock. The artificial CDRH3 sequences are XXXXXXXXXX^1^ and XXXXXXXXXX^1^, while natural CDRH3 sequences are XXXXXXXXXX and XXXXXXXXXX. **c**, Structure of the binding complex formed by PALM-generated CDRH3 XXXXXXXXXX targeting the HR2 region of SARS-CoV-2. The PALM-generated CDRH3 successfully targets the stable peptide region within HR2, which is the input sequence of PALM. **d**, Molecular complex visualization depicting the binding of a natural antibody to the HR1 regions of SARS-CoV-2. It’s noteworthy that the natural antibody does not exhibit binding to the stable peptide in the HR2 region.

We utilized PALM to generate antibodies against the specified antigens in the 14H dataset and then used A2binder for evaluation. It is worth noting that A2binder is a fast screening method that helps us quickly screen for potential high-affinity antibodies. To further validate the effectiveness, we conducted subsequent validation. The results indicated that the affinity prediction values of some of the generated sequences were higher than those of the natural sequences. One CDRH3 region generated by PALM was XXXXXXXXXX^1^, of which the predicted binding free energy is 1.70, smaller than the other generated CDRH3 and the natural CDRH3, XXXXXXXXXX.

To further validate the performance of the PALM, we selected the highest-affinity antibodies from the A2binder’s predictions and conducted structural simulation using AlphaFold2 ^38^. We retrieved the crystal structure of the HR2 region of the SARS-CoV-2 virus from the PDB database ^39^ and used the ClusPro ^40,41^ to perform antigen-antibody docking. For comparison, we also performed the same docking process on the natural antibodies. To investigate the ability of the A2binder, we employed SnugDock ^13^ to adjust the pose of the antigen-antibody complex. It is worth noting that the validation of docking may not be entirely accurate, but it is a widely used computational method for antibody assessment. Docking has aided in the development of numerous antibody design approaches. We employ docking as an external validation tool to further discern the affinity of antibodies selected through A2binder screening.

As shown in Figure 6b, the A2binder predicted that the optimal generated sequence, XXXXXXXXXX^1^, had the lowest binding surface energy and Interface Root Mean Square Deviations (Irmsd), while the natural sequences XXXXXXXXXX and XXXXXXXXXX had higher energies which mean fewer stable bonds. Another artificial sequence, XXXXXXXXXX^1^, which was predicted to have higher affinity than the natural sequence, exhibits lower binding surface energy than the natural sequence by SnugDock. This result validates the predictive ability of A2binder for binding energies.

Figure 6c-d illustrates the results of the docking results between the generated and natural antibodies and the crystal structure of the HR2 region of SARS-CoV-2 (PDB ID: 7ZR2 ^42^). The generated antibody was observed to form hydrogen bonds between the CDRH3 region and the stable peptide in the HR2 region at 101R and 3D sites. In contrast, the natural antibody was found to form hydrogen bonds with the HR1 region instead of the stable peptide in the HR2 region. This demonstrates that PALM is capable of generating antibodies that bind to the correct sites on the SARS-CoV-2 virus. And this capability of targeting the stable regions of the antigen is essential in the design of antibodies against rapidly evolving viruses such as SARS-CoV-2.

### Comparison of PALM with previous methods in antibody design

We used PALM to generate 1000 antibody CDRH3 sequences for the HR2 region of the SARS-CoV-2 virus, which were found to have Levenshtein distances from 4 to 10 from natural antibodies, as shown in Figure 7a. To compare the efficiency of PALM with traditional antibody design methods, we utilized two widely used antibody optimization tools, Rosetta^12^ and Absolute!^43^, to design antibodies for the given epitope of HR2. More specifically, we used the Rosetta SnugDock program for antibody-antigen docking. SnugDock expansions on the standard RosettaDock approach to predict antibody-antigen complexes by optimizing antibody-degrees of freedom relative to binding. The general idea of using Rosetta^12^ and Absolute!^43^ in antibody design is to replace amino acids of natural antibodies and subsequently assess the efficacy of the modified antibody using these tools. Exploring all possible combinations of amino acid changes is impossible due to the computational resources required by these tools. Thus, a popular strategy is to change the amino acid sequentially^23^. This involves changing the amino acid at one position during each design round and assessing the impact on affinity. The altered antibody with the highest affinity is then utilized as the foundation for introducing a new amino acid modification in the next round. Figure 7b presents the CPU hours required for the three methods to generate antibodies at different distances from natural ones. PALM exhibits great advances in saving the computational resources of antibody design. Notably, due to the direct generation of results with different edit distances, PALM’s computational consumption is not affected by an increase in distance. Our results demonstrate that PALM outperforms previous methods in terms of efficiency, which has significant implications for the rapid design of antibody drugs. Such a strategy saves computational resources but has limitations, such as the potential to become trapped in local optima, which hinders exploration of the global fitness landscape of the sequence space. As a result, the traditional strategy may lead to suboptimal or ineffective antibody designs. To investigate whether antibodies designed by PALM exhibit a higher affinity than the traditional strategy, we employed EvoEF2^44^ to design antibodies using the traditional strategy and compared their affinity to those designed by PALM. To enable a fair comparison, we selected the antibody generated by PALM which was deemed optimal by A2binder and had an edit distance of 7. Therefore, we utilized EvoEF2 to perform seven rounds of single-point mutation on natural antibodies and selected the generated antibody with the highest affinity. Figure 7c displays the comparison of the interface energy of antibodies obtained from SnugDock^13^ with the highest affinity generated by PALM and EvoEF2, indicating that the binding surface energy of antibodies generated by PALM is significantly lower than that of EvoEF2. This comparison further emphasizes the advantages of PALM in antibody design.

**Figure 7.**
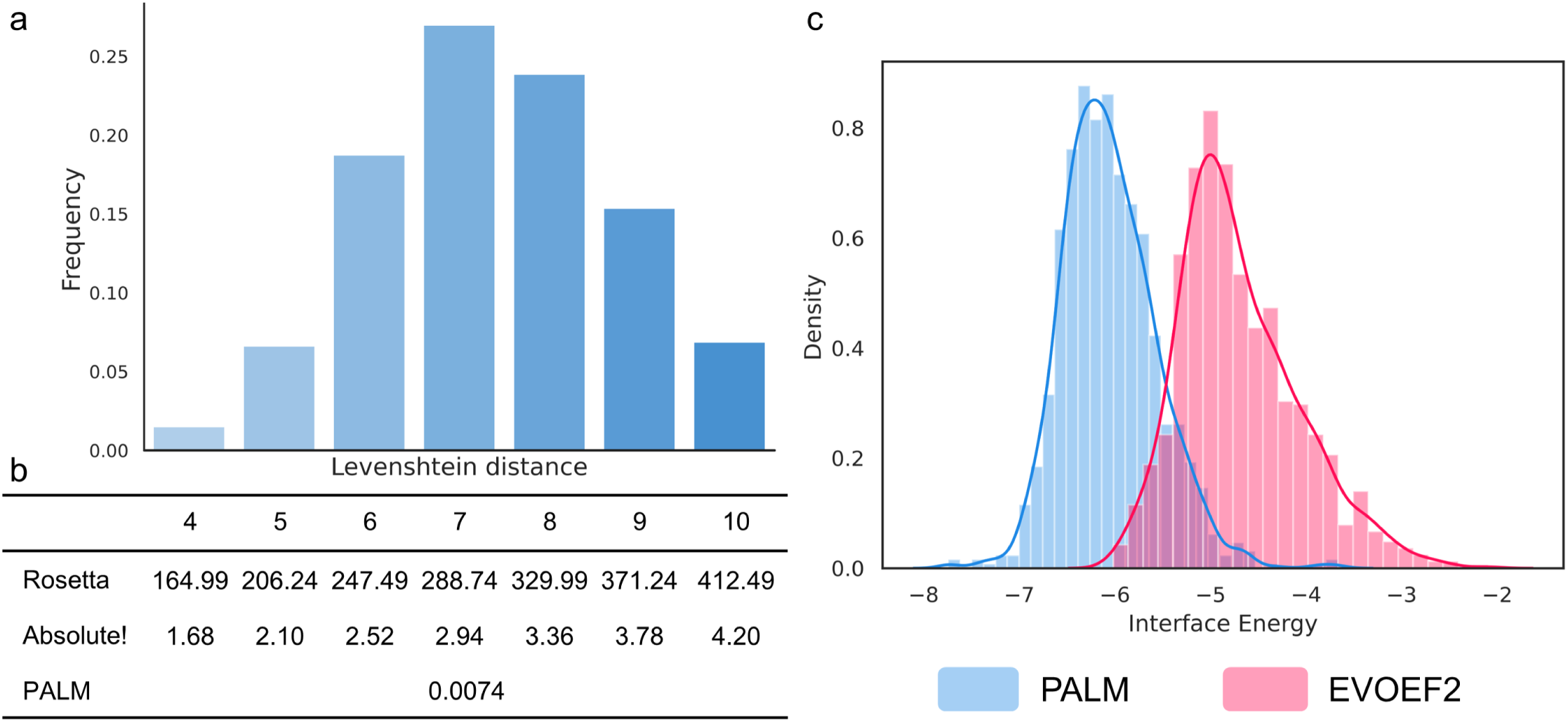
Comparison between PALM and traditional computational antibody design methods. **a**, Distribution plot showcasing the Levenshtein distance among antibodies generated using PALM. **b**, A comparison of the time expenditure for antibody design at varying Levenshtein distances from natural antibodies is conducted among Rosetta, Absolute!, and PALM. The top row illustrates various Levenshtein distances, while the subsequent three rows represent the time required by each method to design antibodies at these distances to natural antibody, measured in CPU hours. **c**, Comparison of the binding affinity, indicated by interface energy, between antibodies produced by PALM and those generated by EvoEF2. The interface energy values were determined independently through SnugDock.

### PALM can generate antibodies that exhibit high binding affinity to novel variants of SARS-CoV-2

To investigate whether the antibodies generated by the PALM exhibit stronger binding abilities against new variants of SARS-CoV-2, we attempted to design high-affinity antibodies targeting these variants using PALM. The CoV-AbDab dataset contains over 20 variants of SARS-CoV-2, but it does not include the new variant XBB, which is rapidly spreading worldwide and rapidly increasing in proportion among all infected cases. Therefore, we aimed to use the PALM to generate specific antibodies against XBB. After training in all positive samples in the CoV-AbDab dataset, we selected the antibodies with the highest predicted binding affinity from the CoV-AbDab dataset and formed a set of similar antibodies for further specific training. We obtained the sequence of the XBB variant from the NCBI database ^45^ and inputted the RDB sequence of XBB into the AF2 ^38^. We replaced the CDRH3 region of the natural antibody targeting to XBB with the generated results and used ClusPro ^40,41^, SnugDock ^13^, and A2binder to evaluate the binding affinity against XBB. One CDRH3 region generated by the PALM was XXXXXXXXXX^1^, and A2binder predicted a degree of neutralization of 3.01, which was higher than the other generated CDRH3 and the natural antibody.

After that, we employed tools such as AF2^38^ and ClusPro^40,41^ to conduct structural validation. To further validate the binding abilities of the generated antibodies, we input the docking results of the generated antibodies and natural antibodies from ClusPro^40,41^ into SnugDock ^13^, a paratope structural optimization algorithm based on RosettaDock ^46^, for structure optimization.

We analyzed the 1000 optimized results of generated and natural antibodies on the Rosetta server and found that the Interface Energy of generated results is significantly lower than that of the natural antibodies (Figure 8a), and the Interface Root Mean Square Deviation (Irmsd) is also generally lower for the generated antibodies (Figure 8b). We assume that the distribution of the artificial results is smaller than the natural results, and the p-value in the independent t-test is 5.9e-48. These findings indicate that the generated antibodies have improved binding capabilities to the antigen. The results suggest that PALM has the potential to generate higher affinity antibody sequences against new variants of SARS-CoV-2.

**Figure 8.**
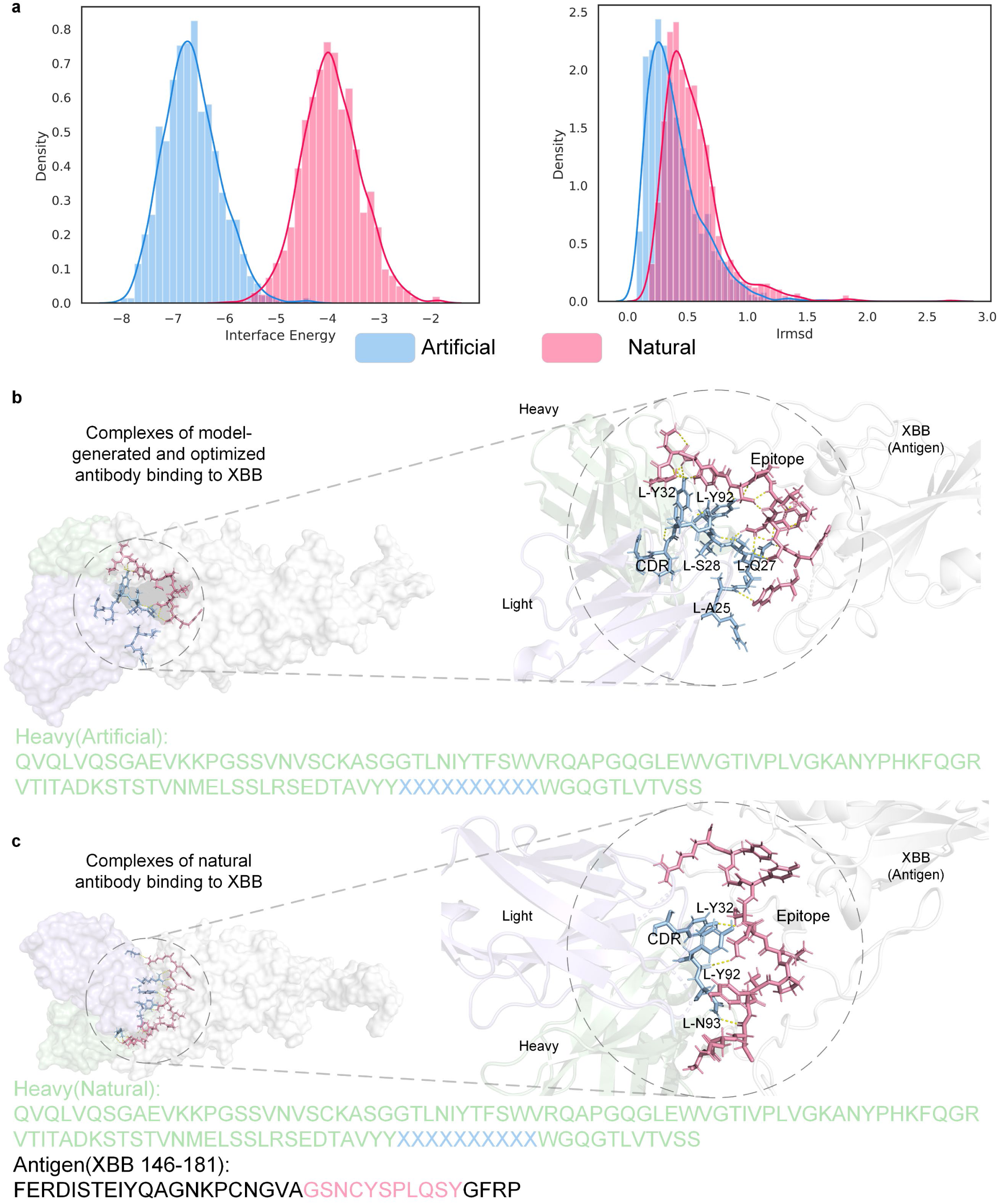
Evaluating PALM’s efficacy in creating artificial antibodies against the novel SARS-CoV-2 variant XBB. **a**, Density distribution plots comparing the affinity of artificial and natural antibodies to XBB, based on 1000 optimized poses of Interface Energy and Irmsd acquired through SnugDock simulations. The artificial antibodies are generated by PALM for XBB, while the natural antibodies are known to target Omicoron variants. Independent t-test results unequivocally demonstrate that both the Interface Energy and Irmsd values of the artificial antibody are significantly lower than those of the natural antibody, underscoring its heightened affinity for variant XBB. **b-c**, Structures of the binding complex formed by the PALM-generated artificial antibody binding to variant XBB. The right side shows a zoomed-in portion of the binding interface on the left, with residues involved in antigen-antibody docking interactions highlighted. The artificial antibody sequence is depicted below the structure, while the PALM-generated CDRH3 sequence is highlighted in blue. The subgraph b corresponds to the left plot of subgraph a, while subgraph c corresponds to the right plot of subgraph a.

Figures 8c and 8d illustrate the docking results and Rosetta optimization results of the generated antibodies and natural antibodies. We can observe that the binding region of the generated antibodies to XBB is more concentrated, primarily binding to the light chain residues A25, Q27, S28, Y32, and Y92. In contrast, the binding sites of the natural antibodies are dispersed and include light chain residues Y32, Y92, and N93.

### PALM is highly interpretable

In addition to its ability to generate high-affinity antibodies against new variants of SARS-CoV-2, we also investigated the interpretability of our model. By utilizing the attention mechanism, PALM may have the potential to focus on key sites during the learning process. We inputted the artificial antibody and the HR2 sequence of SARS-CoV-2 mentioned in section “PALM can generate antibodies that bind to the desired epitopes on the antigens” into PALM. We then computed the average of the multi-layer attention weights of the model.

Figure 9a illustrates the attention weights output by PALM, with red indicating high attention weights and blue indicating low attention weights. The intensity of the color represents the strength of attention. Our analysis revealed that the attention weights of the correct docking sites in PALM’s output were generally high, with the highest attention values observed at the R residues in the CDRH3 region, which forms hydrogen bonds with D residues in the HR2 peptide segment. This suggests that PALM can correctly capture key contact sites, providing insight for further research and optimization of antigen-antibody binding.

**Figure 9.**
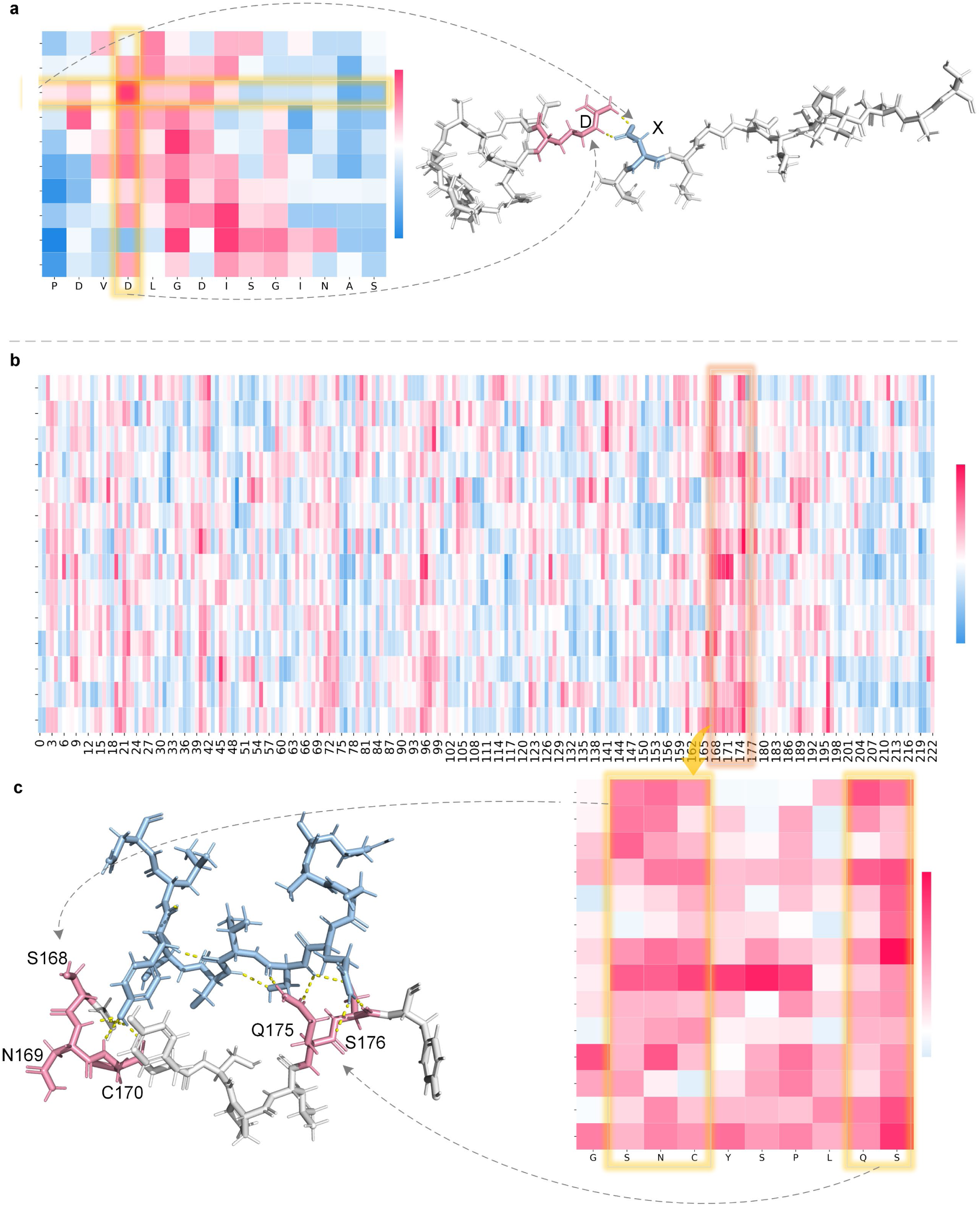
Interpretability analysis of PALM in generating antigen-specific antibody CDRH3 sequence. **a**, Heat maps displaying cross-attention values of PALM when generating CDRH3 sequence “XXXXXXXXXX1” that targets the epitope “PDVDLGDISGINAS” of SARS-CoV-2. Notably, residue D of the epitope and residue R of the CDRH3 region of the antibody exhibits the highest interaction attention values. Cosistent with the cross-attention values, in the binding complexes shown on the right, these two residues form a hydrogen bond link between them. **b**, Heat maps displaying cross-attention values of PALM when generating CDRH3 sequence “XXXXXXXXXX1” that targets the SARS-CoV-2 variant XBB. **c**, Cosistent with the high cross-attention values of the residue 167-177 in the SARS-CoV-2 variant XBB, these reidues play important roles in binding to the generated CDRH3.

Moreover, we analyzed the ability of the model to generate high-affinity antibodies against the new variant XBB. Figure 9b illustrates the attention weights generated by PALM. We observed that the model exhibited a higher attention weight on the region 167-177 of the antigen, specifically corresponding to the binding pocket of XBB and the antibody. Figure 9c shows a zoomed-in view of this region, which indicates that the attention weights are generally higher than the average. Additionally, the key positions for hydrogen bond formation between the antigen and the antibody, S168-C170, and Q175-S176, were found to have high attention values. Among these key positions, only C170 had an attention weight lower than the average, while all other key positions had attention weights higher than the average. We observed that the region 167-177 of the antigen contains XBB-specific mutation sites: S168, N169, and Q175. A previous study has shown that S168 may confer resistance to RBD class 1 and 2 mAbs, while N169 contributes to resistance against RBD class 3 mAbs^47^. Additionally, previous studies have indicated that the Q175 mutation in XBB restores its receptor affinity, thereby restoring its fitness^48,49^. These findings further suggest that the model may be able to correctly identify and capture key positions of antigen-antibody interaction, pointing the direction for further investigation of the XBB variant.

To further validate the model’s interpretability, we performed statistical tests on PDB structures from the BioMap^35^ dataset. Specifically, we used Pymol ^50^ to identify potential hydrogen bond locations between chains, then compared to the mean attention weights from PALM between those chains. This allowed assessing whether the model attends to structurally interacting residue positions. We divided attention weights into two groups – those at hydrogen bond sites versus elsewhere. An independent t-test revealed significantly higher attention at hydrogen bond locations compared to other sites (p<0.01). This provides further evidence that the model captures key interacting positions between antigens and antibodies.

### *In*-*vitro* assays of artificial and natural antibodies

To further validate the effectiveness of antibodies generated by PALM against the wild-type spike protein of SARS-CoV-2, we selected the top-ranked Artificial 1 antibody along with Artificial 2 antibody based on their predicted binding probabilities by A2binder and two natural antibodies, Natural 1 and 2 (Figure 6b). We then evaluated their binding ability using *in-vitro* assays. The Western blot analysis demonstrated that Artificial 1 and 2 were capable of binding to the spike protein at levels similar to or even surpassing, those of natural antibodies (Figure 10a). To further determine their binding affinity and neutralization capability, we conducted surface plasmon resonance analysis and pseudovirus neutralization. Artificial 1 demonstrated high binding affinity with an equilibrium dissociation constant (KD) of 0.05 nm, and superior neutralization potency with a half maximal inhibitory concentration (IC50) of 0.023 μg/ml, compared to all the tested natural antibodies (Figure 10c). The above assays demonstrated that PALM could generate antibodies surpassing natural antibodies for antigens known in training.

**Figure 10.**
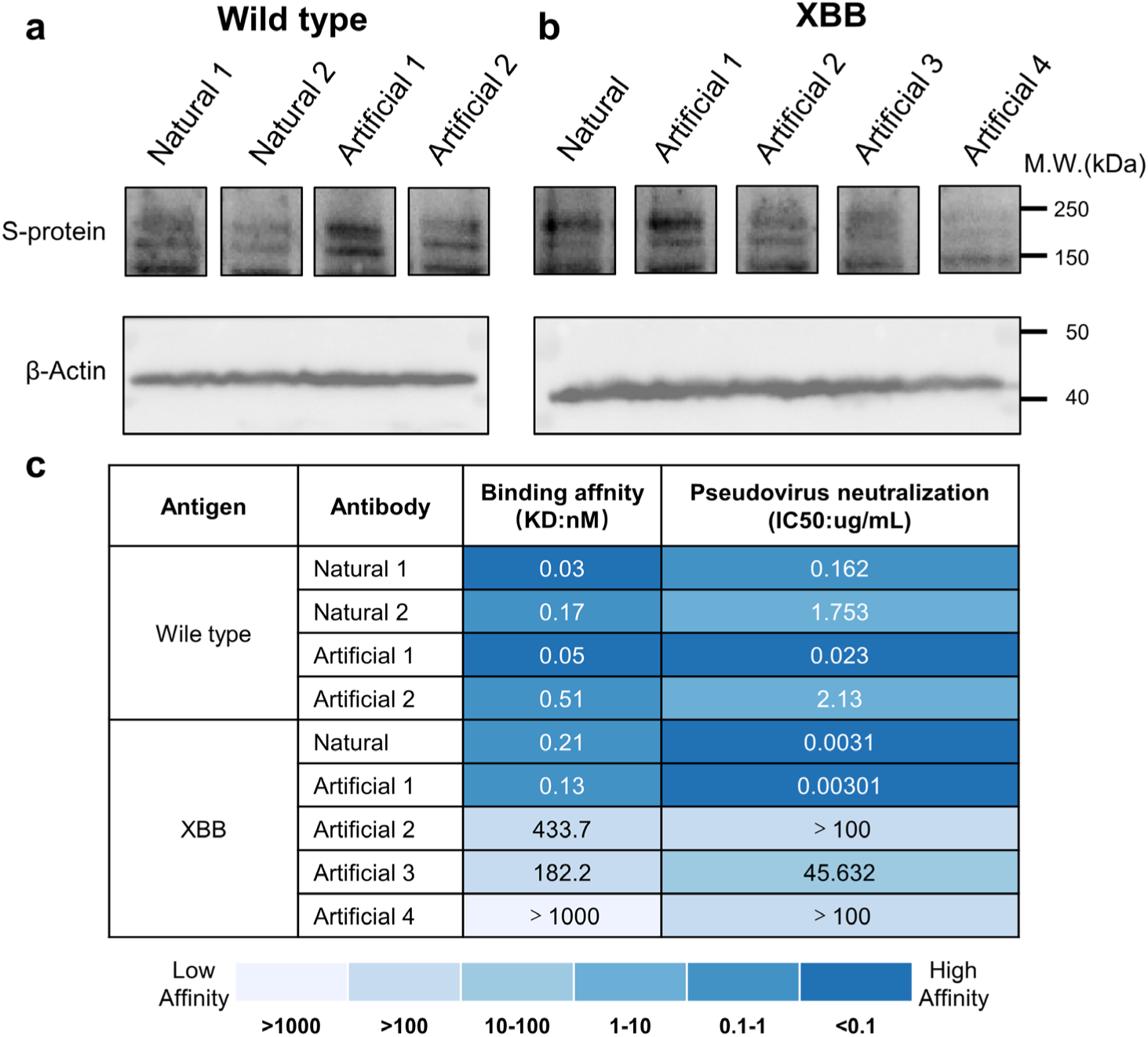
*In-vitro* assays of the binding affinity and neutralization of artificial and natural antibodies. **a-b**, Western blot analysis of artificial and natural antibodies. Antibodies in subfigure **a** target wild-type SARS-CoV-2 S protein, while antibodies in subfigure **b** target SARS-CoV-2 XBB variant S protein. HEK293T cells are used to produce pseudotyped vectors. The x-axis indicates the sample of each band, and the y-axis shows the position of antigen binding. Band intensity demonstrates the affinity between the corresponding antibody and antigen. β-Actin bands at the bottom monitor loading consistency across samples. **c**, The result of surface plasmon resonance analysis and pseudovirus neutralization assays of artificial and natural antibodies. The color legend below indicates value ranges for different colors. Binding affinity and neutralization are measured by KD and IC50, respectively. Lower values signify stronger binding affinity to the antigen and more potent neutralization capability of the antibody.

We next evaluated PALM’s ability to generate artificial antibodies against the novel Omicron variant XBB, which represents a more challenging test case as the model did not see this antigen during training. Like the *in-silico* evaluation shown in Figure 8a, we selected the top-ranked artificial antibody Artificial 1 predicted by A2binder against XBB, along with the natural XBB antibody for the *in-vitro* assays. Besides that, to evaluate the A2binder model using *in-vitro* assays, we also randomly selected three other artificial antibodies, Artificial 2, 3, and 4, which had moderate and lower predicted binding probabilities. Western blot analysis validated the binding of these antibodies to the XBB spike protein (Figure 10b). To further quantify their functional activity, surface plasmon resonance analysis and pseudovirus neutralization assays were performed. As shown in Figure 10c, Artificial 1 demonstrated higher binding affinity, with a KD of 0.13 nm, compared to the natural antibody, and superior neutralization potency against XBB, with an IC50 of 0.00301 μg/ml. The improved performance of Artificial 1 despite no prior exposure to XBB proved PALM’s capacity to generate highly potent antibodies even against novel antigen variants. Consistent with the lower bind probabilities predicted by A2Binder, Artificial 2-4 showed much lower affinities and neutralization than Artificial 1 and natural XBB antibodies. This demonstrated A2binder’s capability to effectively guide the antibody selection for further wet-lab investigations.

## Discussion

In this study, we introduce PALM, a method for generating high-affinity antibody CDRH3 sequences targeting a specific antigen, and A2binder, a method that pairs antigen epitope sequences with antibody sequences to predict the binding specificity and affinity between them. We combined PALM with A2binder to select antibodies with even higher affinity. The efficacy of the A2binder was evaluated by comparing its performance to that of baseline models on various affinity datasets. The results revealed that A2binder exhibited superior antigen-antibody affinity prediction ability, outperforming baseline models on all datasets.

A2binder demonstrates superior performance on affinity datasets, partly attributed to the pre-training of antibody sequences, which enabled the A2binder to learn the unique patterns present in these sequences. The results show that A2binder outperforms the baseline model ESM-F, which has the same framework but the pre-trained model is replaced with ESM2, on all antigen-antibody affinity prediction datasets, suggesting that pre-training with antibody sequences can be beneficial for related downstream tasks.

However, we observed a slight decrease in performance for A2binder and other baseline models on the 14H and 14L datasets compared to other datasets. This observation is consistent with previous studies ^28^. It may be due to a lack of predictive power for large affinity variants arising from only a small number of mutations in the LL-SARS-CoV-2 database, which only contains 1-3 amino acid mutations in antibody sequences.

We explored the differences between the antibodies generated by PALM and natural antibodies. We found that there were significant differences in their sequences, but the binding probability of the generated antibodies was not significantly affected by these differences. Meanwhile, differences in their structures did result in a decrease in binding affinity. These results are consistent with previous studies about network analysis of the antibody repertoires^51^, and functional protein sequences generation^18^. And the potential causes of these phenomena may be due to the binding affinity of an antibody being determined by its ability to recognize and bind to a specific antigen. The structure of its binding site plays a critical role^52^. Small changes in the structure can affect its ability to bind with a high affinity^53–55^. PALM captures important features and patterns contributing to binding affinity through its training on a large dataset of natural antibodies. It can generate antibodies with different sequences while retaining key features necessary for high binding probability. Overall, our results demonstrate that PALM is capable of generating a diverse range of antibody sequences with high binding affinities, despite their dissimilarity to natural antibodies.

Moreover, we validated the performance of PALM by ClusPro^40,41^ and SnugDock^13^, which are widely accepted antibody validation tools^56–59^. PALM was able to generate antibody CDRH3 sequences targeting the stable peptide in the HR2 region of SARS-CoV-2. It generated novel CDRH3 sequences, and the generated sequence XXXXXXXXXX^1^ was validated to demonstrate improved targeting of the stable peptide of the antigen compared to the natural CDHR3 sequence XXXXXXXXXX. Generating antibodies that can accurately target specific antigen sites is crucial in developing effective disease treatment plans ^60^. Rapid and precise antibody generation could lead to the development of new and more effective therapies for various diseases ^61^. Furthermore, PALM was able to generate antibody CDRH3 sequences with higher affinity against the newly emerged variant XBB of SARS-CoV-2. The generated sequence XXXXXXXXXX^1^ displayed a stronger affinity to XBB than natural CDHR3 sequence XXXXXXXXXX. PALM’s output encompasses multiple amino acid variations in CDRH3, surpassing the capabilities of previous antibody optimization models that only allowed for single-point mutations.

In addition, we conducted the *in-vitro* assays, including Western blot, surface plasmon resonance analysis, and pseudovirus neutralization assays, providing critical validation of the efficacy of PALM-designed antibodies. Both the antibodies targeting the spike protein of SARS-CoV-2 wild-type and XBB variant generated by PALM achieved superior binding affinity and neutralization potency compared to natural antibodies in these assays. The strong empirical results from these wet-lab experiments complement the computational predictions and analyses, providing validation of PALM’s and A2binder’s capabilities in generating and selecting robust antibodies with high specificity and affinity against both known and novel antigens.

Our validations on antibodies against the new SARS-CoV-2 variant XBB demonstrate the potential of using a pre-trained model to rapidly generate high-affinity antibodies against a new variant of the virus. This result is significant as it provides a promising approach to developing effective treatments for emerging and evolving viruses. Additionally, this approach could potentially be applied to other infectious diseases, allowing for more efficient and targeted development of treatments.

Unlike the previous antibody optimization methods, we used the pre-training model to learn the construction mode of antibody sequence and then employed the transformer-like model architecture to directly generate the sequence from antigen to antibody key region, which may accelerate the process of antibody design.

At the same time, we used a multi-scale feature fusion neural network to extract global features of the pre-training model. It improves the ability of the A2binder to predict the affinity and assists PALM in finding antibodies with higher affinity.

Significant limitations remain within our methods and more broadly within the study of antibody design. The first is the few minimally curated antigen-antibody pair datasets that exist currently. Due to the lack of antigen-antibody pairing data, we did not directly pre-train the encoder-decoder model, instead opting to train the models for antigen and antibody separately. Also, this led us to generate only CDRH3. Furthermore, our work as a general framework still lacks design for universal antibodies, but rather experiments and validation for SARS-CoV-2 antigens with abundant antigen-antibody pairs. Besides, the validation of antibodies using computational methods such as A2Binder and SnugDock docking methods still has limitations. Docking methods can exhibit instability when calculating binding energies. Additionally, while the high-affinity CDRH3 sequences generated by PALM are promising, certain motifs like the RR dipeptide and Trp residue in XXXXXXXXXX^1^ may increase the risks of polyreactivity and rapid clearance ^62–64^. Such developability liabilities could make these sequences less suitable as clinical candidates despite high predicted affinity. Future work should explore approaches to balance affinity with minimized developability risks, such as filtering certain motifs or optimizing to remove liabilities like the RR and Trp while maintaining potency. Advanced models could also be trained to directly predict developability profiles. There is a substantial need to better integrate proper antibody draggability into computational design. In the coming future, we aim to procure more comprehensive antigen-antibody data from richer sources to enable the creation of universal antibody bi-chains. Furthermore, we aim to explore the possibility of further engineering on this molecule, aiming to preserve its affinity while mitigating any associated drawbacks. Nonetheless, our study serves as evidence that generative models hold great potential in producing high-affinity antibodies.

In conclusion, the proposed PALM integrates the capability of large-scale antibody pre-training and the effectiveness of global feature fusion, resulting in superior affinity prediction performance and the ability to design a high-affinity antibody. Furthermore, the direct sequence generation and the interpretable weight visualization make it an efficient and insightful tool for designing high-affinity antibodies.

## Methods

### Datasets

Observed Antibody Space (OAS): The OAS database ^31,32^ compiles and annotates an extensive collection of immune repertoires, encompassing over one billion sequence data from more than 80 studies. These data encompass a broad spectrum of immune states and subjects. A significant discrepancy exists between the volume of unpaired sequence data and paired data within this dataset. We isolated sequences from the unpaired data set that corresponded to the human species and obtained from individuals diagnosed with SARS-CoV-2 at the time of sequencing. This resulted in a selection of 81,750,886 unpaired heavy chains and 17,754,502 unpaired light chains.

The Coronavirus Antibody Database (Cov-AbDab): Cov-AbDab^34^ is a public database that catalogs all published and patented antibodies and nanobodies with the capability to bind to coronaviruses, including SARS-CoV-2, SARS-CoV-1, and MERS-CoV. It comprises 11,868 studies, each of which encompasses multiple data pairs that indicate the neutralizing capacity of the antibody towards a specific virus strain. During the curation process, we restricted our focus to the 10,720 studies that included both the light and heavy chain sequences of the antibodies, resulting in a total of 35,970 antigen-antibody pairs. Subsequently, we removed data with indeterminate locus positions. Finally, we retained only the data of the variants associated with the SARS-CoV-2 antigen, which resulted in a total of 27,324 unique data entries.

14H and 14L: The 14H and 14L datasets are sourced from the LL-SARS-CoV-2 database^36^. This database was designed to choose two heavy chains and two light chains from three main chains of antibodies as the framework for constructing an antibody library. Each dataset includes 1-3 mutations in the CDR region of the skeleton sequence. The AlphaSeq^65^ technology was applied to quantify the affinity binding of each sequence variant to the SARS-CoV-2 target. As part of the data processing, we eliminated raw data that lacked affinity information for both 14H and 14L datasets. Then, we averaged the affinity measurements for all sequences with the same antibody sequence. This resulted in 13,922 unique heavy chain data entries in 14H and 18,708 unique light chain data entries in 14L. In the 14H dataset, only the heavy chain is varied, with the light chain and antigen remaining the same for each entry. Similarly, in the 14L dataset, only the light chain is different, while the heavy chain and antigen remain constant for all entries.

BioMap^35^: BioMap data set was derived from an antigen-antibody affinity prediction competition held by BioMap. It contains 1,706 antigen-antibody pairing data and has binding free energy (Delta G) as the label. The antigen-antibody sequences and Delta G data in the BioMap dataset are sourced from the Biomap company, specifically from 473 PDB complex entries. It comprises 638 unique antigens and 1,277 unique antibodies, with most antibodies of human and mouse origin along with minor fractions from hamsters, chimpanzees, rhesus monkeys, rabbits, rats, and llamas. No nanobodies are included.

### Rotary Transformer (RoFormer)

Roformer improves the Transformer structure using Rotary Position Embedding (RoPE). This design achieves relative position coding in the form of absolute position coding. Experimental results on various long text classification benchmark datasets show that the enhanced Transformer with rotary position embedding, namely RoFormer, can give better performance compared to baseline alternatives and thus demonstrates the efficacy of the proposed RoPE. Suppose 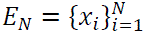 is the word embedding of N input tokens. Self-attention first merges the location information into the word embeddings and converts them into q, k, and v representations:

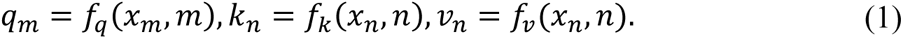

The information of the *m*^*th*^ and *n*^*th*^ positions are obtained by functions *f*_*q*_, *f*_*k*_ and *f*_*v*_. The attention weights are then obtained by q and k:

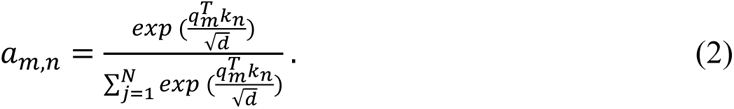

As seen in equation (2), 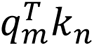 can transfer knowledge between tokens at different positions. To merge relative location information, RoPE requires that the inner product of the query *a*_*m*_ and the key *k*_*n*_ is represented by a function *g* that takes as input variables only the word embeddings *x*_*m*_, *x*_*n*_ and their relative locations *m* − *n*:

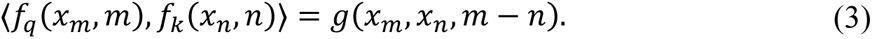

Su et al.^21^ assumed that the inner product encodes position information only in the relative form. After derivation, they obtained the representation of RoPE:

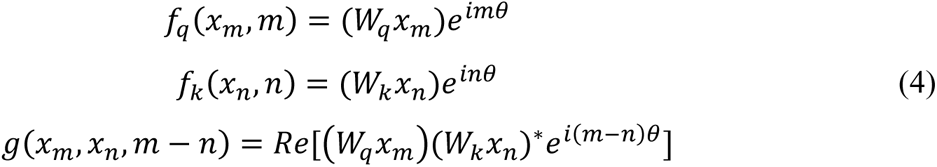

Where (*W*_*k*_*x*_*n*_)^∗^ represents the conjugate complex number of *W*_*k*_ *x*_*n*_, and *Re*[·] is the real part of a complex number, θ ∈ *R* is a preset non-zero constant. Details of the derivation of the above equation can be found in the original paper^21^.

The RoPE rotates the affine-transformed word embedding vector by a specific angle multiple of its position index. It employs absolute position encoding to achieve relative position encoding and eliminates the need to operate the Attention matrix. As a result, RoPE has the potential to be used with linear attention. Considering the conformation of amino acid folding in proteins, relative position-coding may be better suited for constructing linkage patterns between proteins, such as antigens and antibodies, when modeling a language model.

### MF-CNN

We use an elaborate multi-scale feature fusion CNN architecture MF-CNN to further fuse the sequence features extracted by Roformer.

The input of MF-CNN is the output of all the tokens of Roformer. It utilizes a 3-layer CNN skeleton containing convolution, pooling, and Relu for multi-scale feature extraction.

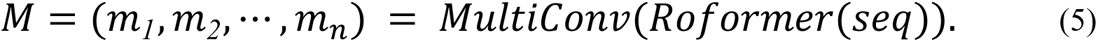

The features are then further fused using a multi-layer FC layer and residual operations for the final output.

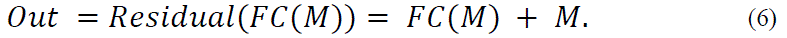

### Baseline

ESM-F: ESM-F is based on the ESM2 ^16^. It modifies pre-trained Roformers of A2binder to ESM2s and retains the MF-CNN model for feature fusion. ESM-2^16^ is a state-of-the-art protein representation model. It is suitable for fine-tuning a wide range of tasks that take protein sequences as input. They used a BERT-style encoder-only transformer architecture with modifications. They changed the number of layers, number of attention heads, hidden size, and feed-forward hidden size as they scaled the ESM model. The purpose of constructing ESM-F as a baseline is to investigate the extent of improvement that can be achieved in the antigen-antibody affinity model by employing pre-training with antibody sequences.

Ens-Grad: The Ens-Grad model is derived from the high-capacity CNN architecture proposed by Liu et al.^66^ for antibody CDR design. Each input token of the architecture is converted into a one-hot vector, where each position of the vector is the channel input of CNN. All sequences are filled with zero to the maximum sequence length of the dataset. The model is composed of two convolutional layers followed by a standard FC decision layer. Using Ens-Grad as the baseline allows us to compare the performance of A2binder and the antigen-antibody affinity prediction model without pre-training.

Vanilla BERT: The BERT is based on Transformer’s encoder architecture^30^. It consists of multiple encoder layers, and each layer contains self-attention and feed-forward modules. It assigns token position information through absolute position coding. We selected Vanilla BERT as the baseline to investigate the impact of RoPE, MF-CNN, and pre-training on the model’s performance.

### Pre-training

We employed the Unique Amino Acid (UAA) Tokenizer during pre-training and downstream fine-tuning, as antibodies are highly responsive to single-point mutations. The UAA Tokenizer works by assigning a unique token to each unique amino acid present in the protein sequence. Each token represents a specific amino acid, which allows models to understand and analyze the sequence. The UAA tokenizer assigns each amino acid a unique integer value. After UAA tokenization, a weighted, learnable embedding layer converts each amino acid’s integer value into a dense vector representation. This embedding allows the model to learn the complex features of each residue about its position. This retains positional relationships between residues, enabling the model to better learn mutations, deletions, and insertions.

Roformer’s model parameters refer to BERT-light. The hidden size is 768, including 12 hidden layers and 12 attention heads. The intermediate size is 1,536.

MAA task is a self-supervised pre-training method. Through a part of the token of the random mask input sequence, it uses the model to predict the mask position and conduct self-supervised learning of the model. We randomly masked 15% of the tokens to conduct MAA training.

For the pre-training of heavy chain Roformer, we used 81,730,448 training data and 20,438 test data, for the pre-training of light chain Roformer. 17,736,747 training data and 17,755 test data were used.

The batch size of the pre-training was set to 2048. The heavy chain model was trained for more than 200,000 steps, while the light chain model was trained for more than 100,000 steps.

Both heavy and light Roformer were trained on NVIDIA Volta V100 GPUs using a distributed computing architecture.

### Antigen-antibody Affinity Prediction

In the antigen-antibody affinity prediction task, we split the training, validation, and test sets in a ratio of 8:1:1. To improve the model’s performance, we employed two learning rates for optimizing the Roformer and MF-CNN, respectively. Additionally, a linear learning rate schedule was implemented to facilitate an effective training process. We repeat experiments five times by choosing different random seeds and reporting the average results for our model and baseline methods.

In the binding specificity classification task carried out on the Cov-AbDab dataset, we used the ROC-AUC, PR-AUC, and Accuracy as evaluation metrics. In the affinity prediction task, we used Spearman’s rank correlation coefficient as metric.

### Antibody Generation

Since the antibodies in the 14H dataset target a stable peptide in the HR2 region of SARS-CoV-2, to train the cross-attention layer in the Antibody Layer of the decoder, we first utilized positive samples from the CoV-AbDab dataset to train PALM for 10 epochs. In the case of generating the correct target key sites, we used the antigen-CDRH3 paired data from the 14H data set for further fine-tuning. In the case of antibody generation for the new variant XBB, we selected positive samples with the same 15 amino acids of light and heavy chain prefixes in the antibody for further training.

The Beam search method was used in the generation phase. Beam search can reduce memory consumption through a breadth-first searching strategy, which is widely employed to boost the output text quality. It often leads to substantial improvement over the greedy search strategy, which is equivalent to setting the beam size to 1. We set the Beam size to 10. After generation, we replaced the CDRH3 of the natural sample, used A2binder trained on the corresponding dataset for affinity prediction, and selected the best of them for subsequent validation of the results.

### Western blot

The antigen protein was separated by SDS-PAGE and transferred to an NC membrane. After blocking, the membrane was incubated with different antibodies overnight at 4°C. Then the membrane was incubated with a secondary antibody conjugated to HRP for 1 hour at room temperature. ECL substrate was applied to visualize antibody binding bands. The Western blot results were imaged using a chemiluminescence imaging system.

### Surface plasmon resonance analysis

The interaction between RBD and antibodies was analyzed by surface plasmon resonance using a BIAcore T200 instrument. PBS running buffer containing 0.005% Tween 20 was used. The RBD analyte was injected into the PBS buffer for buffer exchange. Antibodies were captured on the sensor chip, and then serial dilutions of RBD flowed over the surface to obtain binding data. Curve fitting was performed to calculate each antibody-RBD pair’s dissociation constant (KD).

### Pseudovirus neutralization assay

Pseudoviruses expressing spike proteins were produced in HEK 293T cells and harvested from cell culture supernatant. The pseudoviruses were incubated with serial dilutions of antibodies starting from 100 μg/mL for 1 hour at 37°C to allow neutralization. The antibody-pseudovirus mixtures were then added to Vero cells seeded in 96-well plates to infect the cells. After overnight incubation, pseudovirus transduction units were quantified by imaging cytometry to generate neutralization curves. The half-maximal inhibitory concentration (IC50) was determined for each antibody.

## Code and data Availability

Pre-trained Roformer and well-trained PALM and A2binder models on the comprehensive training dataset are available on Zenodo: https://doi.org/10.5281/zenodo.7794583. The source code of PALM are also available at: https://github.com/TencentAILabHealthcare/PALM.

## Author Contributions

H.H: Methodology, Software, Formal analysis, Writing – original draft, Visualization. B.H.: Conceptualization, Supervision, Methodology, Investigation, Data Curation, Writing – Original Draft. L.G.: Wet Lab Validation. Y.Z.: Software, Resources, Data Curation. G.C.: Visualization, Formal analysis. Q. Z.: Wet Lab Validation. C.Y.-C.C.: Supervision, Writing – Review & Editing. T. L.: Supervision, Wet Lab Validation. J.Y.: Conceptualization, Supervision, Writing – Review & Editing.

## Acknowledgements

This work was supported by the National Natural Science Foundation of China (Grant No. 62176272), Research and Development Program of Guangzhou Science and Technology Bureau (No. 2023B01J1016), and Key-Area Research and Development Program of Guangdong Province (No. 2020B1111100001).

## Competing interests

The authors declare no competing interests.

## Footnotes

1 under patent application

## Notes

https://github.com/TencentAILabHealthcare/PALM

### Competing Interest Statement

The authors have declared no competing interest.

